# The mechanism of checkpoint-dependent DNA replication fork stabilization in human cells

**DOI:** 10.1101/2024.11.01.621514

**Authors:** Agostina P. Bertolin, Berta Canal, Mona Yekezare, Jingkun Zeng, Rachael Instrell, Michael Howell, John F.X. Diffley

## Abstract

The DNA damage checkpoint is crucial for maintaining genome stability after genotoxic stress; without it, excess DNA replication origins are activated, stalled DNA replication forks cannot restart normally, high levels of DNA damage and single-stranded DNA (ssDNA) accumulate and cells cannot complete S phase. Preventing excess origin firing suppresses all these effects. Here we show that when replication is not restrained by a functional checkpoint, excess DNA synthesis sequesters the processivity factor PCNA and its loader RFC, preventing normal fork restart. Nascent DNA ends unprotected by RFC/PCNA are attacked by the Helicase-Like Transcription Factor (HLTF), causing irreversible replication fork collapse and hyperaccumulation of single-stranded DNA. This explains how the checkpoint stabilizes stalled replication forks and has implications for how origin firing is normally coordinated with fork progression. Loss of HLTF suppresses fork collapse and cell lethality in checkpoint-deficient cells, which has implications for how resistance to anti-checkpoint therapies may arise.

## Introduction

DNA replication fork stalling in response to nucleotide depletion, inhibition of DNA polymerases or DNA damage activates the checkpoint protein kinases ATR and its downstream effector, Chk1^1–3^. In budding yeast, mice, and humans, these damage checkpoint kinases are essential for cell viability^1,4–12^ and checkpoint defects in humans are associated with inherited developmental diseases, premature ageing, and predisposition to various types of cancer^13–18^. These kinases coordinate a wide range of cellular responses to promote genome stability and survival^19^, including regulation of damage-dependent transcription, DNA repair, and cell cycle arrest; however, the DNA replication-associated functions of the checkpoint appear to be especially important^20–22^. The checkpoint inhibits replication origin firing in yeast and humans; in yeast by inhibiting two replication origin firing factors, Sld3 and Dbf4^23,24^ and in humans by targeting Cdc25A, the Cdk2 phosphatase and activator, for ubiquitin-mediated proteolysis^25–27^. The checkpoint also protects stalled replication forks in yeast and humans from undergoing catastrophic changes often referred to as ‘fork collapse’^28^. Replisomes stall irreversibly and can no longer resume replication, even if the checkpoint is restored^29,30^, and normal DNA replication fork structures are converted into structures associated with recombination^30^, despite the fact that CMG helicase and associated factors remains associated with DNA^31^. How checkpoints stabilize replication forks and the mechanistic basis for irreversible fork arrest are currently poorly understood. Nucleases including Exo1^32–34^, MRX^35^ and the proofreading exonuclease of DNA polymerase ε^36^ have all been implicated in this phenomenon in yeast. Evidence from human cells indicates that deregulation of origin firing in the absence of a functional checkpoint drives fork collapse^28^ and that accumulation of ssDNA during this process depletes the soluble pool of the ssDNA binding protein RPA, which causes DNA damage^37^. In this manuscript we have re-examined events occurring at stalled replication forks in the absence of a functional checkpoint in human cells and we provide evidence that depletion of Replication Factor C (RFC), the PCNA sliding clamp loader, by excess Okazaki fragments that is the initial driver of fork collapse, and that the Snf2-family DNA translocase HLTF executes the irreversible fork collapse.

## Results

### Checkpoint inhibition after replication fork stalling induces S phase catastrophe

Consistent with previous work showing that loss of ATR or Chk1 causes a diverse array of pathological phenotypes^1,11,17,37–41^, treatment of U2OS cells with ATR or Chk1 inhibitors (ATRi, Chk1i) caused cell lethality (**Figures S1A** and **S1B**) which was exacerbated by variety of genotoxic agents (**Figures 1A** and **S1C**-**S1F**), including the DNA polymerase α, δ, and ε inhibitor aphidicolin (aph), which we use extensively in this manuscript. Addition of Chk1i during the last 4h of a 24h aphidicolin treatment increased DNA damage markers, chromatin-bound Rad51, increased ssDNA derived from both parental and nascent DNA and caused RPA exhaustion as evidenced by cessation of RPA accumulation on chromatin followed by accumulation of γH2AX (**Figures S1G**-**S1K**) consistent with previous work^38–40,42^. Cells released from this treatment regime (aph/Chk1i, **Figure S1L**) were largely unable to complete S phase and enter mitosis, in contrast to cells released from aph-only, and the few cells that reached mitosis exhibited high levels of nuclear aberration (**Figures 1B**, **1C** and **S1M**).

**Figure 1.**
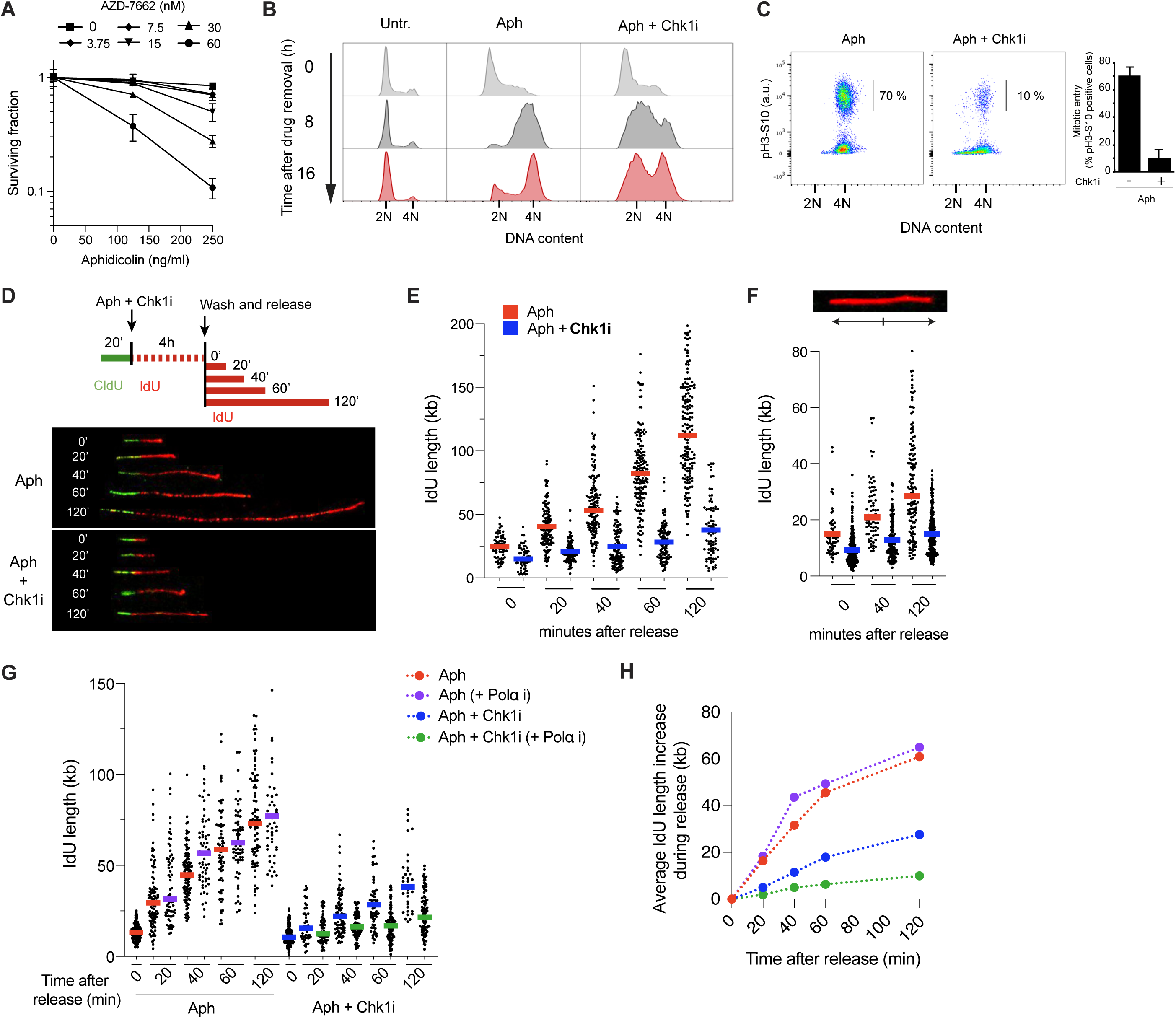
Checkpoint inhibition after replication fork stalling induces S phase catastrophe. **(A)** Survival assay of U2OS cells treated with indicated doses of Chk1i (AZD7762) in combination with aphidicolin. Viability was determined using CellTiter-Glo and normalised to aph-untreated cells. **(B)** S phase progression analysis. 0, 8 and 16h after release in media with 100 nM taxol. **(C)** Mitotic entry analysis. 16h after release in media with 100 nM taxol. The proportion of pH3-S10 positive cells is shown. **(D)** DNA fiber analysis of individual replication fork progression. Upper panel, experimental setup: cells were pulsed with CldU for 20 min, washed and incubated with IdU and aph alone or in combination with Chk1i for 4h. Cells were then washed and incubated again only with IdU for 20-, 40-, 60- and 120-min. Bottom panel, representative micrographs of CldU-IdU replication tracks. **(E-F)** Scatter plot and mean IdU track length in bicolour fibers (**E**) or red-only fibers (**F**). **(G)** DNA fiber analysis of Polα-dependent synthesis following checkpoint inhibition. Experimental setup as in **D** but the Polα inhibitor ST1926 (4 μM) was added during release. Scatter plot and mean IdU track length in bicolour fibers. **(H)** Average length increase for IdU tracks of bicolour fibers during **release.**

Chk1 inhibition also had a profound effect on DNA replication fork restart (**Figures 1D**-**1F**), consistent with previous work^38,40,43–45^. Cells were pulsed with CldU (green) for 20min, CldU was washed out, and aphidicolin and IdU (red), with or without checkpoint inhibitors, were added. After 4 hours, aphidicolin and IdU were washed out and IdU alone was added back (**Figure 1D**). Replication forks from cells released from aph-only treatment immediately began to synthesise DNA after aph removal at rates very similar to unperturbed forks (**Figures 1D** and **1E**). Replication forks also resumed synthesis after aph/Chk1i treatment, however, fork rate was roughly 5 times slower (**Figures 1D** and **1E**). As expected^43,44,46–48^, transient checkpoint inhibition led to a significant increase in new origin firing measured as tracks labelled only in red (**Figure S1N**). These red-only tracks were much shorter after Chk1i (**Figure 1F**) consistent with the idea that replication forks are affected by Chk1i regardless of whether they were established before or after checkpoint inhibitor addition. We did not detect nascent strand degradation^49^ in either aph- or aph/Chk1i-treated cells (**Figure S1O**) indicating that the reduction in apparent extension rates observed after fork stalling in checkpoint deficient cells is a consequence of reduced replication fork elongation rate, not increased degradation rate, consistent with previous work^38,50^.

Fork rate when assessed by DNA fiber spreading was not affected when the Polα inhibitor, ST-1926^51,52^, was added during the restart of replication forks in aph-only cells; however, ST-1926 almost completely inhibited fork extension in aph/Chk1i-treated cells (**Figures 1G** and **1H**). Similar results were obtained with a second Polα inhibitor, CD-437^53^ (**Figure S1P**). Although DNA fiber spreading cannot distinguish leading from lagging strands^54^, the synthesis of many kilobases of DNA after aph alone in the presence of the Polα inhibitor is most consistent with efficient resumption of leading strand replication, probably in the absence of lagging strand synthesis; the near complete inhibition of all synthesis after aphidicolin and Chk1i treatment can only be interpreted as loss of both leading and lagging strand synthesis. Therefore, the slow DNA synthesis during replication fork restart in checkpoint deficient cells is dependent upon Polα whilst the rapid synthesis seen in active forks after release from aph-only does not require Polα and is more consistent with resumption of processive leading strand DNA synthesis by Polδ and/or Polε.

### Order of effects following checkpoint inhibition

We next determined the order that the myriad phenotypes first appeared after checkpoint inhibition. We exposed aph-treated cells to increasing durations of Chk1 inhibitor, from 10 min to 5 hours (**Figure S1Q**). DNA damage markers were almost undetectable until 45 min after checkpoint inhibitor addition (**Figure 2A**) while RPA exhaustion, defined as the point where γH2AX levels continue to rise without commensurate increase in chromatin-bound RPA, was first seen after 60 min of treatment and continued to accumulate over the next 4 hours (**Figure 2B**). A clear increase in origin firing could be detected even at the earliest time points (10-20 min) after checkpoint inhibitor addition when compared to aphidicolin alone (**Figures 2C**, **2D** and **S1R**) consistent with previous work^38^. Defective replication fork restart was also observed early, after 20 min of Chk1i treatment (**Figure 2E**). These results show that new origin firing and, importantly, fork defects occur well before the onset of detectable DNA damage or depletion of chromatin-bound RPA arguing that DNA damage generation or RPA exhaustion are likely to be downstream of new origin firing and/or fork defects following checkpoint inhibition.

**Figure 2.**
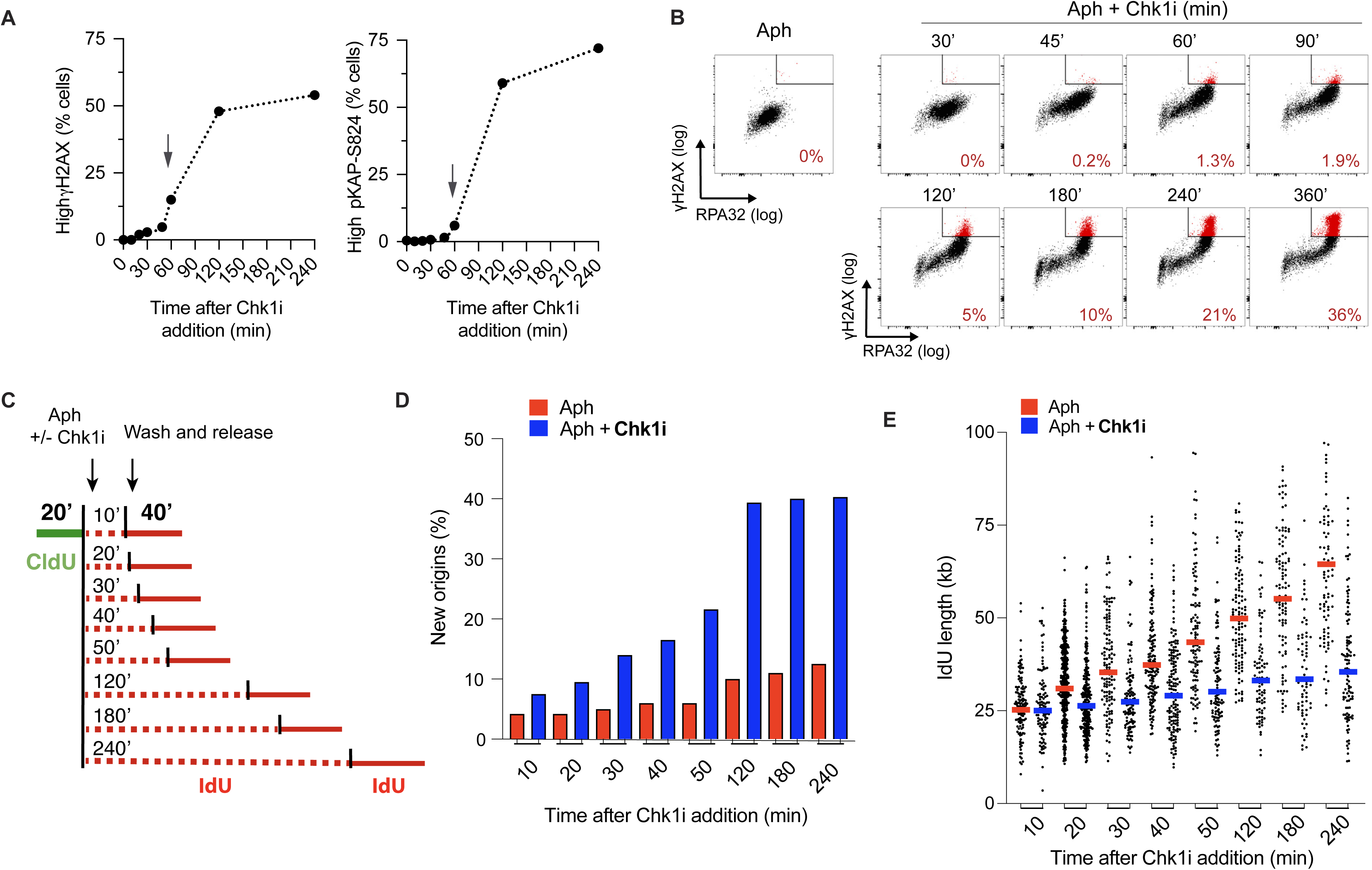
Order of effects following checkpoint inhibition. **(A-B)** Flow cytometry analysis of DNA damage markers as depicted in **Figure S2A**. **(A)** Percentage of cells presenting high γH2AX and phospho-S824 KAP1 levels in cells treated with aph alone or in combination with Chk1i. **(B)** Levels of chromatin-bound RPA32 and γH2AX in the same population of cells. Representative images comparing γH2AX intensity (log scale) and chromatin-bound RPA32 intensity (log scale) in cells. Gated cells depicted in red represent cells with simultaneously high levels of γH2AX and RPA. Bottom right of each panel, percentage of red coloured cells. **(C-E)** DNA fiber analysis of replication fork progression and origin firing. **(C)** Experimental setup: cells were pulsed with CldU for 20 min, washed and then incubated with IdU plus 1 μg/ml aph alone or in combination with 50 nM Chk1i for the indicated time. Cells were then washed and released in complete media containing IdU for 40 min. **(D)** Percentage of new origins fired was determined by counting the number of IdU-only tracts over the total number of different replication structures. **(E)** Scatter plot and mean IdU track length in bicolour fibers.

### Excess origin firing depletes chromatin-bound PCNA and causes replication fork defects

New origin firing and fork restart defects started at similar times after Chk1i treatment. Inhibition of excess origin firing using either Cdc7 or Cdk2 inhibitors suppressed DNA damage markers, RPA exhaustion and the defect in replication fork restart associated with Chk1i (**Figures 3A**-**3C** and **S2A**), in agreement with previous work^37,44^ and also rescued mitotic entry and viability (**Figures S2B** and **3D**). This suggests that the generation of an excessive number of replication forks depletes some essential replication factor. It has previously been proposed that this limiting factor is RPA: treatment with hydroxyurea and checkpoint inhibitors generates sufficient ssDNA to exhaust the soluble pool of RPA, resulting in DNA damage, fork breakage and ultimately fork collapse^37^. However, our results indicate that fork defects precede RPA exhaustion. RPA overexpression (**Figure S2C)** rescued some of the DNA damage seen at 120 min and later (**Figure S2D**) consistent with previous work^37^. However, it did not rescue the DNA damage observed at earlier time points. Moreover, RPA overexpression did not rescue the Chk1i-induced fork restart defects at any time point (**Figure S2F**). Consistent with this, chromatin-bound RPA levels are relatively low in the first 30 min when restart defects first occur, but continue to rise steadily over time, and do not plateau until 4 hours of checkpoint inhibition (**Figure 3E**). Thus, RPA is unlikely to be the replication factor limiting for fork restart seen from early time points, though its depletion does contribute to DNA damage at later time points.

**Figure 3.**
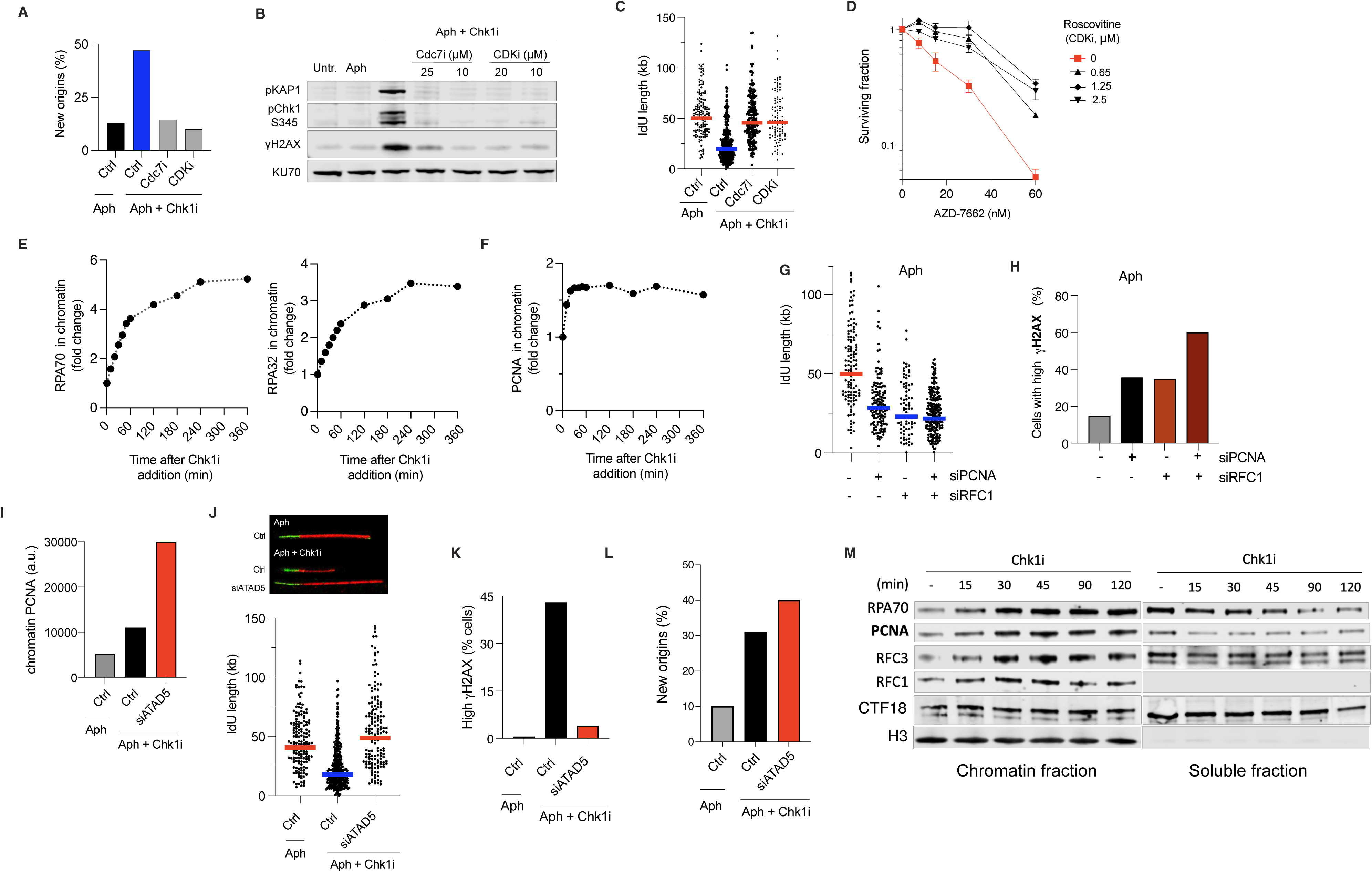
Excess origin firing depletes chromatin-bound PCNA and causes replication fork defects. **(A-C)** Cells were treated with aph alone or in combination with Chk1i with and without Cdc7i (XL-143) or CDK2i (roscovitine). **(A)** Percentage of new origins fired. **(B)** DNA damage markers immunoblots. **(C)** Scatter plot and mean IdU track length in bicolour fibers. **(D)** Survival assay of cells treated with indicated concentrations of Chk1i alone or in combination with roscovitine. **(E-F)** Chromatin-bound RPA32 and RPA70 (**E**) and PCNA (**F**) in aph/Chk1i-treated cells. Graphs show fold change of mean fluorescence intensities (MFI) in comparison with aph alone. **(G)** Replication fork progression by DNA fiber analysis in aph-treated cells depleted for PCNA and RFC1 alone or in combination. Scatter plot and mean IdU track length in bicolour fibers. **(H)** Percentage of cells presenting high levels of γH2AX in aph-treated cells depleted for PCNA and RFC1 alone or in combination. **(I)** Immunoblot of chromatin extracts of aphidicolin treated cells for 24h alone or with the addition of 50 nM Chk1 inhibitor during the indicated times just before harvesting. Histone 3 (H3) was used as a loading control for chromatin fraction. **(J)** MFI of chromatin-bound PCNA in ATAD5 depleted cells. **(K)** DNA fiber analysis of replication fork progression in ATAD5 depleted cells. Left, scatter plot and mean IdU track length in bicolour fibers. Right, representative micrographs. **(L)** Percentage of control or ATAD5 depleted cells presenting high levels of γH2AX. **(M)** Percentage of new origins fired in control and ATAD5 depleted cells.

In sharp contrast to RPA, however, we found that the amount of PCNA bound to chromatin reached a maximum level within 30 min of checkpoint inhibition (**Figure 3F**) despite the fact that the number of new forks continued to rise over 2 hours (**Figures 2D** and **S1R**). We therefore considered this decrease in the amount of PCNA per fork could be responsible for the replication fork restart defect, analogous to results in the accompanying manuscript. Consistent with this and with previous work^55^, transient reduction of either PCNA or the PCNA clamp loader RFC1 (**Figure S2G)** caused DNA damage and affected replication restart, even in the absence of excess origin firing and regardless of checkpoint status (compare **Figures 3G** with **3H** and with **S2H**)^55^, suggesting that reduction of chromatin-bound PCNA levels prevents normal fork restart. Transient overexpression of PCNA led to a significant increase of DNA damage markers in aphidicolin conditions alone (**Figures S2I** and **S2J**), suggesting non-specific effects. We therefore turned to a different approach, depletion of the main PCNA unloader, ATAD5/Elg1^56,57^. ATAD5 was among the top 10 hits in a whole-genome CRISPR-Cas9 dropout screen in RPE1 cells for genes whose loss conferred resistance to the ATR inhibitor AZD6738^58^, ranking higher than known sensitivity determinants such as Cdc25A and Cdc25B^59^. ATAD5 was also identified as a top hit in a similar whole-genome CRISPR-Cas9 dropout screen for AZD6738 in mouse embryonic stem cells^60^. ATAD5 depletion led to a significant increase in chromatin-bound PCNA (**Figure 3I**). This almost completely suppressed the Chk1i-induced fork restart defects and accumulation of DNA damage markers (**Figures 3J** and **3K**). Co-depletion of PCNA, but not Rad18, with ATAD5 prevented suppression of DNA damage (**Figures S2K** and **S2L**), arguing that suppression is due to the increased chromatin-bound PCNA and not to an increase in K164 mono-ubiquitylation of PCNA. Importantly, ATAD5 depletion did not prevent excess origin firing (**Figure 3L**) consistent with previous work^61^.

PCNA is an abundant protein, and, consistent with this, even after 120 min of Chk1 inhibitor exposure, PCNA, as well as RPA, were detected in soluble as well as chromatin fractions (**Figure 3M**). Similarly, RFC3, one of the core RFC subunits present in all RFC complexes, was also detected in both fractions. By contrast, RFC1, which is roughly 10-fold less abundant than the core RFC subunits and ∼20 fold less abundant than PCNA^62^, was detectable on chromatin but not in the soluble fraction, even at the earliest times (**Figure 3M**). This argues that, during excess of origin firing, RFC1-5, rather than PCNA is depleted.

### HLTF is responsible for the defects caused by RFC/PCNA-depleted forks in the absence of the checkpoint

Our results thus far show that excess origin firing drives RFC/PCNA depletion, which prevents normal replication restart. They do not, however, explain where the DNA damage and excessive ssDNA come from or why the fork arrest is irreversible. We developed an RNA interference-based genome-wide screen in human cells transiently treated with aph and Chk1 inhibitors where we monitored γH2AX levels as a marker for DNA damage and the mitotic phosphorylation of H3 (pH3-S10) as an indicator of S/G2 phase completion (for further details see **Figure S3** and **Tables S1** and **S2**). When focusing on the siRNAs that increased γH2AX levels, we found that out of the ∼21,000 genes tested, RFC2 depletion emerged as the highest-scoring gene, RFC1 depletion ranked among the top 20, and all RFC subunits increased γH2AX levels (**Figure S3E**), further supporting the idea that RFC1-5 is the limiting factor preventing normal fork restart.

Next, we identified genes whose knockdown reduced DNA damage and promoted mitotic entry after Chk1i removal, indicating S-phase completion through fork resumption. After secondary screens and siRNA deconvolution, one of the strongest such suppressors was Helicase-Like Transcription Factor (HLTF, also known as SMARCA3) (**Figure S3F** and **Table S2**). HLTF is a fork-associated protein, ubiquitin ligase and SNF2-family DNA translocase implicated in several DNA repair/tolerance pathways^63–68^. Moreover, HLTF inactivation reduces DNA damage and slightly increases resistance to genotoxic agents and ATR inhibitors^64^. We found that both HLTF siRNA depletion and CRISPR/Cas9 gene knockout (HLTF-KO) strongly reduced phosphorylation of all DDR markers analysed in aph-treated cells after Chk1 or ATR inhibition (**Figures 4A**, **S4A** and **S4B**). Complementation of HLTF-KO cells with full-length HLTF cDNA (**Figure S4C)** restored the elevated DNA damage induced by checkpoint inhibition (**Figure S4D**). High levels of parental and nascent ssDNA, Rad51 foci and DSBs were almost completely suppressed in aph/Chk1i cells in the absence of HLTF (**Figures 4B** and **S4E**-**S4G**). This reduction in S phase-related DNA damage in HLTF deficient cells was accompanied by enhanced levels of mitotic entry and a strong reduction in Chk1i-induced genomic instability (**Figures 4C** and **S4H**) consistent with these cells being able to resume replication and complete S phase (**Figure S4I**). HLTF depletion and gene knockout suppressed the replication restart defects in ATR- or Chk1-inhibited cells from replication forks established either before or after Chk1 inhibition (**Figures 4D**, **4E**, **S4J** and **S4K**). Notably, the suppression of restart defects seen in HLTF-deficient cells occurred despite the excess origin firing and depletion of chromatin-bound PCNA (**Figures S4L**-**S4N**). Consistent with this, HLTF ranked among the top 5 hits in CRISPR dropout screens for resistance to ATR and Chk1 inhibitors using a DNA damage sgRNA sub-library^69^. Moreover, it scored among the top hits in whole-genome CRISPR-Cas9 screens in MCF10^70^ and RPE1^58^ cells for resistance to the ATR inhibitor AZD6738.

**Figure 4.**
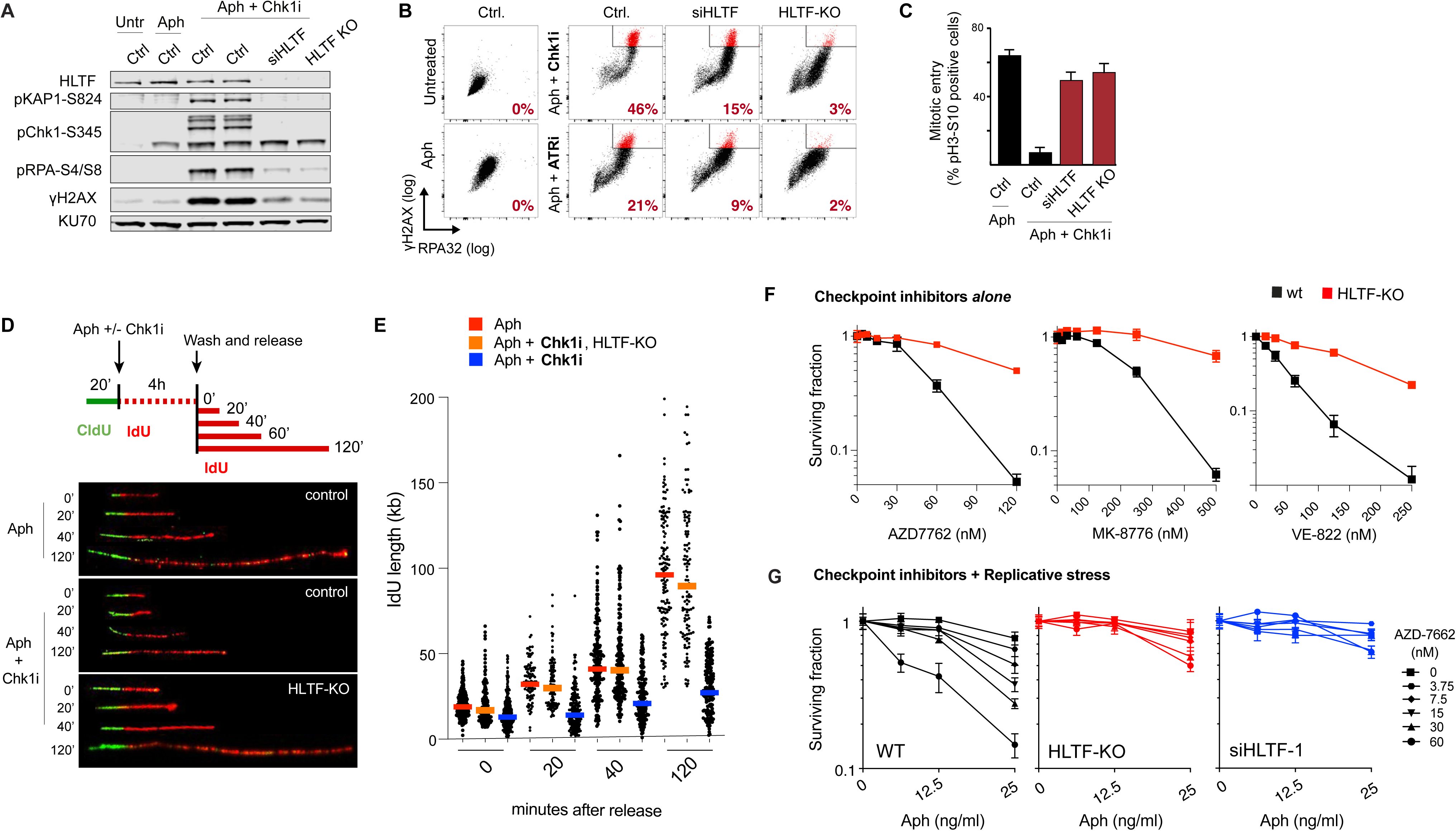
HLTF is responsible for the defects caused by RFC/PCNA-depleted forks in the absence of the checkpoint. **(A)** Immunoblots for DNA damage markers in control, HLTF siRNA knock-down (KD) and HLTF CRISPR knock-out (KO) cells after checkpoint inhibition. **(B)** Flow cytometry analysis of chromatin-bound RPA32 and γH2AX levels in the same population of cells. Gated cells depicted in red represent cells with simultaneously high levels of γH2AX and RPA. Bottom right of each panel, percentage of red coloured cells. **(C)** Mitotic entry analysis. Cells were treated with aph for 24h, with or without Chk1i during the final 4 hours. Cells were washed and released for 16h. The proportion of pH3-S10 positive cells is shown. **(D, E)** Kinetic analysis of replication fork progression and origin firing in checkpoint inhibited conditions by DNA fiber. **(D)** Upper panel, experimental setup. Bottom panel, representative micrographs. **(E)** Scatter plot and mean IdU track length in bicolour fibers. **(F, G)** Survival assay of HLTF-KO or KD cells treated with indicated concentrations of Chk1i (AZD7762, MK-8776) or ATRi (VE-822) alone (**F**) or in combination with aph (**G**).

To further investigate the drivers of fork collapse, we examined cellular processes involving HLTF. HLTF, along with the E2 ubiquitin-conjugating complex Mms2/Ubc13, polyubiquitylates PCNA in response to DNA damage to initiate error-free lesion bypass via template switching^66,71,72^. We generated MMS2 and Ubc13 CRISPR-KO cells, neither of which suppressed DNA damage in the absence of checkpoint activity (**Figure S5A**), suggesting PCNA polyubiquitination is not involved in fork collapse. SHPRH, like HLTF, is a human Rad5 ortholog with a RING domain and SWI/SNF-ATPase helicase but lacks the HIRAN domain^67,72,73^. We generated SHPRH KO cells, and they show levels of DNA damage marker phosphorylation similar to controls after checkpoint inhibition, suggesting no involvement in this process (**Figure S5B**). Genetic and biochemical studies suggest that replication fork reversal^74^ is predominantly catalyzed by the SNF2-family DNA translocases HLTF^63,64^, SMARCAL1^75^ and ZRANB3^76,77^ and SMARCAL1 has been shown to be required for the generation of nascent-strand ssDNA in human cells when ATR is inactivated^38^. So, we tested whether SMARCAL1 or ZRANB3 are required to promote fork collapse in the absence of a functional checkpoint. Individual and double CRISPR-KO U2OS cell lines of SMARCAL1 and ZRANB3 did not suppress Chk1 inhibition-induced phosphorylation of DNA damage markers (**Figure S5C**), generation of parental ssDNA or accumulation of RPA and Rad51 foci (**Figure S5D**). Moreover, epistasis experiments revealed that HLTF depletion suppressed Chk1 inhibition-induced DNA damage markers, regardless of ZRANB3 or SMARCAL1 status (**Figure S5E**). These findings suggest that SMARCAL1 and ZRANB3 are not involved in triggering checkpoint inhibition-induced fork collapse.

Our results suggest that depletion of RFC/PCNA by excess OkF synthesis from the excess origins generates a substrate for HLTF to act upon, leading to fork collapse. In line with this, ATAD5 and HLTF depletion alleviated the effects observed when PCNA and/or RFC1 are depleted without excess origin firing and regardless of checkpoint status (**Figures 5A**-**5C**). This suggests that PCNA retention on chromatin can protect free 3’-ends from nuclease attack and that HLTF can cause fork restart defects even in the presence of a functional checkpoint. Consistent with this, overexpressing HLTF without fork stalling or checkpoint inhibition led to an increase in DNA damage markers (**Figure 5D**). These results suggest that the checkpoint may protect replication forks from HLTF even in the absence of genotoxic stress. HLTF-deficient cells showed increased resistance to ATR and Chk1 inhibitors in the absence (**Figure 4F** and **S6F**) and presence (**Figures 4G** and **S6**) of a variety of genotoxic drugs (aph/HU/Cisplatin/Olaparib).

**Figure 5.**
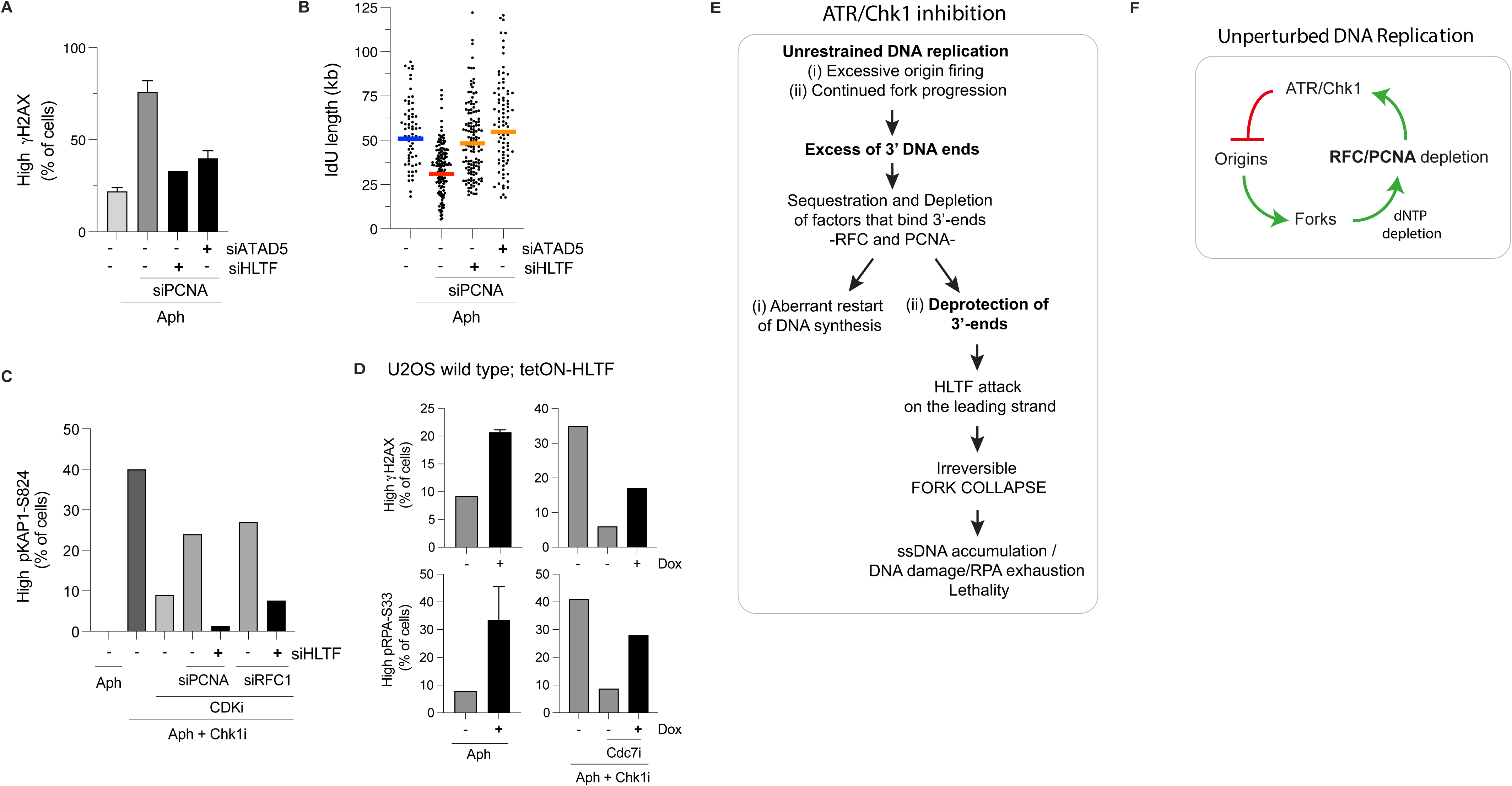
HLTF is responsible for the defects caused by RFC/PCNA-depleted forks. **(A)** Flow cytometry analysis of the percentage of aph-treated cells presenting high levels of γH2AX chromatin staining in control cells and PCNA siRNA treated cells alone or in combination with HLTF or ATAD5 siRNA. **(B)** DNA fiber analysis of replication fork progression in aph conditions in control cells and PCNA siRNA treated cells alone or in combination with HLTF or ATAD5 siRNA. Cells were pulsed with CldU for 20 min, washed and then incubated with IdU plus 1 μg/ml aph for 4h. Cells were then washed and incubated again only with IdU for 40 min. Scatter plot and mean IdU track length in bicolour fibers. At least 120 forks were scored per sample. **(C)** Flow cytometry analysis of the percentage of cells presenting high levels of pKAP1 staining. Control cells were treated with 1 μg/ml aph for 24h alone. Control cells and cells treated with PCNA or RFC1 siRNA alone or in combination with HLTF siRNA were subjected to aph for 24h and a checkpoint inhibitor with or without a CDK2i (roscovitine, 5 μM) was added during the final 4 hrs. **(D)** Flow cytometry analysis of the percentage of tetON-HLTF wild type U2OS cells presenting high levels of DNA damage markers (γH2AX and pRPA-S33) after treatment with aph alone or in combination with Chk11i with or without Cdc7i (XL-143, 10 μM). **(E)** Proposed order of events and dependencies during replication fork collapse in the absence of a functional checkpoint. **(F)** Model for coordination of fork progression and origin firing via checkpoint activation and dNTP/RFC/PCNA depletion.

## Discussion

The work described here, together with accompanying manuscript showing the detailed molecular mechanism through biochemical reconstitution of DNA replication fork stalling and restart, provides a novel framework for understanding how the DNA damage checkpoint stabilizes stalled replication forks in human cells (**Figure 5E**). When DNA replication origin firing is not restrained by the checkpoint after inhibition of DNA synthesis, excess Okazaki fragments (OkFs) are initiated; these OkFs sequester PCNA and the canonical Rfc1-containing RFC complex, which becomes specifically depleted. As incomplete OkFs accumulate (OkF synthesis cannot complete because of DNA polymerase inhibition, see accompanying manuscript), the depletion of RFC/PCNA exposes 3’ DNA ends and it is the action of HLTF on these exposed ends that causes fork collapse (**Figures 5E**). HLTF localises to replication forks even in the absence of genotoxic stress^63^ and promotes fork reversal *in vivo* and *in vitro* via a HIRAN domain that binds to 3’ DNA ends^63,64,78,79^. We propose that the irreversible effect of HLTF on replication forks is the initiation of unwinding of the nascent leading strand^80^, which exposes ssDNA from both the nascent and parental strands (**Figure S4E**) and depletes RPA (**Figure 4B**) consistent with previous work^37,38,40,42^. By eliminating the nascent leading strand, any continued unwinding by CMG after restart can only support lagging strand synthesis, which explains why all synthesis after aph/Chk1i treatment is slow and is sensitive to Polα inhibitors (**Figures 1G** and **1H** and **S1P**). Although PRIMPOL may be able to generate primers on the leading strand template^81^, these cannot be efficiently extended under conditions of RFC/PCNA depletion.

The work in this manuscript emphasises the critical role for checkpoint regulation of origin firing; we do not know whether checkpoint-dependent phosphorylation of Claspin (human orthologue of Mrc1), Mcm10 or other replisome components also contributes to inhibiting OkF synthesis by slowing forks in humans as it does in yeast^82^ (see accompanying manuscript).

Our work also provides insights into the role of the checkpoint in coordinating replication during an unperturbed S phase. HLTF promotes cell lethality after checkpoint inhibition even *without* added replicative stress (**Figure 4F**), HLTF is responsible for the DNA damage induced by partial PCNA or RFC knockdown even in the presence of a functional checkpoint (**Figures 5A**-**5C**) and HLTF overexpression causes DNA damage even in the presence of a functional checkpoint (**Figure 5D**). These results suggest that the balance between HLTF, RFC/PCNA/3’-ends and the DNA damage checkpoint is crucial even during an unperturbed S phase (**Figure 5F**). We suggest that, upon entry into S phase, origin activation continues until the number of forks begins to deplete RFC. RFC depletion causes uncoupling of leading strand synthesis from CMG progression and prevents OkF completion and maturation, generating ssDNA on the leading and lagging strand templates which drive checkpoint activation. Excess origin activation also cause dNTP depletion^41^, which, as shown in the accompanying manuscript, can promote RFC/PCNA depletion and thus may also contribute to checkpoint activation. Checkpoint activation then inhibits new origin firing, preventing further RFC/PCNA depletion; as OkFs mature and replicons terminate, RFC and PCNA are released and recycled and as RFC depletion is relieved, the checkpoint signal is quenched, allowing new origins to fire. Thus, RFC/PCNA/3’-ends homeostasis, reinforced by dNTP depletion, ensures a constant level of origin firing throughout S phase (**Figure 5F**). Consistent with this model, Chk1 is active throughout unperturbed S phase^83^, and inhibition of the checkpoint during unperturbed S phase leads rapidly to increased origin firing^38,43,44,48,84^, indicating that the checkpoint is constantly suppressing new origin firing. Conditions that limit origin activation should prevent RFC depletion and accelerate fork rate and consistent with this, previous work has shown that reducing origin firing by manipulating the levels of Cdc7, CDK, or Orc1 results in faster unperturbed fork rates, supporting this view^85,86^.

HLTF is frequently silenced by promoter hypermethylation in colon, gastric, oesophageal and other cancers^78^ and HLTF knockout promotes malignant transformation in mice^87^. Additionally, tumour cells express high levels of PCNA^88^ and RFC^89,90^ presumably to accommodate their high degree of uncontrolled replication and to overcome RFC/PCNA depletion. This is also consistent with *Atad5* heterozygous mice being cancer prone^91,92^ and with *Atad5* being identified as a genetic risk locus in genome-wide association studies for breast cancer^93^ and serous ovarian cancer^94^. This suggests HLTF functions as a tumour suppressor and that maintaining RFC/PCNA/3’-end homeostasis may serve as a tumour suppressive mechanism. We hypothesise that loss of HLTF may make some cancer cells – perhaps those with compromised DNA checkpoints or with dysregulated origin firing– resistant to chemotherapies. This may be especially important as a route for the emergence of drug resistance in cancers treated with DNA checkpoint inhibitors.

## Materials and Methods

### Cell Culture

The following cell lines used in this study (U2OS, hTERT-RPE1, HeLa Kyoto) were originally obtained from ATCC and were cultured in DMEM (Sigma). eHAP cells obtained from Horizon Discovery were cultured in IMDM medium (Life Technologies). Cultures were supplemented with 10% FBS (Invitrogen), L-glutamine (GlutaMAX, ThermoFisher) and penicillin and streptomycin (Gibco). All cells were grown at 37 °C in 5% CO2. The cell lines used in our study were authenticated by STR profiling in the Crick Cell Service Crick facility. All cells are routinely tested to be mycoplasma free.

### RNA Interference, drugs, and small molecule inhibitors

Knockdown studies were performed using siRNA from Dharmacon. siRNAs were transfected at 5-50 nM final concentration depending on the experiment using Lipofectamine RNAiMAX (Invitrogen) and OptiMEM (Invitrogen). Experiments were completed 2 days after siRNA transfection. The following siRNAs were used: siRNA Control (siGENOME, non-targeting siRNA Pool #2, D-001206-14-20); siRNA HLTF-1 (ON-TARGETplus, SMARTpool, L-006448-00); siRNA HLTF-2 (siGENOME, D-006448-04); siRNA PCNA (ON-TARGETplus, SMARTpool, L-003289-00); siRNA RFC1 (ON-TARGETplus, SMARTpool, L-009290-00). The following used small molecule inhibitors/drugs were used: AZD7762 and MK-8776 (CHK1 inhibitors, 50-100 nM, Axon MedChem), VE-821 and VE-822 (ATR inhibitors, 500 nM, SelleckChem). Zeocin and blasticidin (4 µg/ml) were obtained from Life Technologies. Phalloidin (ThermoFisher). Aphidicolin (0.5-1 mg/ml), nocodazole (100 ng/ml), BrdU (20 µM) and cytochalasin B (4.5 μg/ml), PARPi (Olaparib), cisplatin, hydroxyurea and doxycycline were obtained from Sigma-Aldrich. Cdc7 inhibitor (XL-143, 5 to 10 μM, Axon), CDK2/1 inhibitor (roscovitine, 5 to 10 μM, Calbiochem), DNA-polymerase α inhibitors (CD437, Sigma-Aldrich and ST1926, MedChem).

### Cell lines

*CRISPR-Cas9 Knock-out cell lines*. For creation of HLTF knockout cells, the following gRNA sequences designed using Benchling software based on human HLTF Exon 2 sequence (NM_003071.3): 5’ CACCGTTGGACTACGCTATTACAC 3’ and 5’ aaacGTGTAATAGCGTAGTCCAAC 3’. In all cases, U2OS or eHAP parental cells were transfected using Lipofectamine 3000 (Invitrogen) with pX459-sgRNA (gRNAs targeting unique sequences within each gene locus) and selected briefly with Puromycin for two days. Live single cells were sorted by flow cytometry, plated into 96-well plates, and allow to grow for 2 weeks. Individual clones were screened for the absence of HLTF protein expression by immunoblotting with the specific antibodies (ab17984, Abcam). <u>HLTF over-expressing cell</u> <u>lines.</u> Stable TetON-HLTF cell lines were constructed by random plasmid integration. Human coding sequences of HLTF were cloned using KpnI/NotI restriction enzymes into the pcDNA4/TO vector (Invitrogen) using the following primers:

Forward (KpnI) 5’GG<u>GGTACC</u>ATGTCCTGGATGTTCAAGAGGG 3’
Reverse (NotI) 5’ATAGTTTA<u>GCGGCCGC</u>TTATAAGTCAATTAATGTTC 3’

T-REx-U2OS cells were obtained from Invitrogen. T-REx-HLTF-KO cells stably expressing the Tet repressor were constructed by transfecting HLTF-KO U2OS cells with the pcDNA6/TR plasmid (Invitrogen) and cells were selected in medium containing 5 μg/ml Blasticidin. Live single cells were sorted by flow cytometry, plated into 96-well plates, and allow to grow for 2 weeks. Individual clones were screened for the Tet repressor protein expression by immunoblotting with the specific antibodies (TET02, MoBiTec) and positive clones were selected. T-REx-wild type cells or T-REx-HLTF-KO cells were transfected with Lipofectamine 3000 (Invitrogen) using the pcDNA4/TO constructs carrying human HLTF sequence. Transformed cells were selected in 250 μg/ml Zeocin. Live single cells were sorted by flow cytometry, plated into 96-well plates, and allow to grow for 2 weeks. Individual clones were screened for Doxycycline dependent protein expression by immunoblotting with HLTF antibodies (ab17984, Abcam) and positive clones were selected.

### Flow cytometry and cell sorting

Multiplexed flow cytometry analysis using fluorescent cell barcoding, combined with EdU, antibody and DNA staining, was performed as previously described ^94^. Up to 6 samples subjected to different treatments were barcoded in each experiment to allow unbiased subsequent antibody staining of the combined samples. For detection of EdU, cells pulsed with 10 mM EdU for the indicated times were harvested and stained with Click-iT chemistry using Click-iT EdU Alexa Fluor 647 Flow Cytometry Assay Kit (Thermo, C10424) according to the manufacturer’s instructions. To quantify the amount of proteins in the insoluble/chromatin fraction, cells were treated with cytoskeleton buffer (CSK) 0.25% Triton X-100 for 5 min on ice. For DNA content analysis, cells were treated with 100 mg/mL RNase A and stained with 1 mg/mL DAPI. Data were analysed using FlowJo software. Cell doublets were excluded for all analyses. See Antibodies for details.

### Antibodies

The antibodies used for western blotting were the following: HLTF (ab17984, Abcam); pH2AX-S139 (γH2AX, 1:2000, clone JBW301, Millipore), pChk1-S345 (1:3000, 2348L, Cell signalling); pKAP1 S824 (1:1000, A300-767A, Bethyl); pRPA2-S4S8 (1:7500, A300-245A, Bethyl); KU70 (1:3000, sc-5309, Santa Cruz); alpha-Tubulin (1:4000, Sigma, T5168); PCNA (1:1000, PC10, Santa Cruz); RFC3 (1:1000, ab182143, Abcam); RFC1 (1:1000, ab229229, Abcam); RFC1 (1:1000, sc-271656, Santa Cruz); H3 (1:1000, ab1791, Abcam). Blots were imaged using the LI-COR platform (LI-COR Biosciences) with anti-rabbit RDye 800CW and anti-mouse IRDye680RD secondary antibodies (both 1:10000). Images were analyzed and quantified using the Odyssey Infrared imaging system (LI-COR Biosciences).

The antibodies used for flow cytometry were the following: pH3-S10 (1:200, ab14955, Abcam); pH2AX-S139 (γH2AX, 1:250, clone JBW301, Millipore); PCNA (1:250, PC10, Santa Cruz); RPA32 (1:250, ab2175, Abcam); RPA70 (1:250, EPR3472, Abcam). For chromatin RPA exhaustion/depletion, pH2AX-S139 (γH2AX, 1:250, ab11174, Abcam) in combination with RPA32 (1:250, ab2175, Abcam). The antibodies used for immunofluorescence studies were the following: RPA (Ab-2 NA18, Calbiochem), RPA (9H8 ab2175, Abcam); Rad51 (H92, Santa Cruz); BrdU (Becton Dickinson).

### DNA fiber stretching assay

U2OS cells were pulse labelled with 25 μM CldU for 20 min, washed with PBS and subsequently pulse labelled with 250 μM IdU together with aphidicolin (1 mg/ml) with or without Chk1 inhibitor (AZD7662, 50nM) or ATR inhibitor (VE-821, 500 nM) for the indicated time (from 10 min to 4h). Cells were washed with warm PBS 3 times and immediately harvest or release in complete media with 250 μM IdU for the indicated time (20 to 120 min). Cells were then trypsinised and resuspended at a concentration of 4×10^5^ cell/ml in cold PBS. 3 ul of cell suspension were spotted on clean glass slides, air-dried for 5 min and lysed with 8 ul of 0.5% SDS in 200 mM Tris-HCL, pH 7.4, 50 mM EDTA (2-5 min, RT). Slides were slightly tilted, allowing the drop to run down the slide, air-dried for 10 min and then fixed in fresh methanol/acetic acid (3:1) for 10 min at RT. Then, air dried ON at 4 °C. Samples were then denatured (2.5 M HCl for 75 min). Slides were blocked in 1% BSA/PBS + 0.1% Tween-20 for 30 min at RT and incubated with rat anti-BrdU monoclonal antibody (αCldU; AbD Serotec) and mouse anti-BrdU monoclonal antibody (αIdU; Becton Dickinson), both 1:500 for 1h at RT. Later, slides were incubated with anti-rat IgG Alexa Fluor 488 (Green, Molecular Probes) and anti-mouse and IgG AlexaFluor 555 (Red, Molecular Probes) both at 1:500 for 1.5h at RT. Slides were washed and mounted with Fluoroshield (Sigma). Images were acquired using a Zeiss Upright 710 confocal microscope with 60x or 43x oil immersion objective. Fiber length was measured using FIJI. The pixel size values were converted into micrometres (μm) using the scale bars created by the microscope and then converted to kilobases according to the conversion factor 1 μm = 2.59 kb. For IdU length measures, otherwise indicated, only bicolour fibers were counted (green-red). For new origin firing quantification, origins fired during the second pulse (IdU, red structures) were quantified as percentage of all structures containing red (green-red and red only). Between 200-700 total replication structures were scored in each sample. Experiments were performed in duplicate or in triplicate.

### Nuclear aberration analysis

Transfected cells were replated at low density. 24 hr after replating cytochalasin B (4.5 μg/ml Sigma) was added to the media and 40 hr later cells were washed 1 min with hypotonic buffer (KCl 0.0075 M) diluted 1/10 from stock solution in PBS 1X, twice with PBS 1X and fixed with PFA/sucrose 2% for 20 min. Phalloidin and DAPI staining were used to visualize the whole cell and its nuclei respectively.

### Survival Assay

For colony formation assays, cells were seeded in duplicates (6-cm dishes) per condition and allowed to adhere for a minimum of 6 h. Cells were then treated with indicated concentrations of Aph for 24 h, washed extensively and incubated in drug-free medium for 11 days. Plates were then washed once in PBS, left to dry, and stained with cell staining solution (0.5% w/v crystal violet, 25% v/v methanol). Finally, the plates were washed three times in deionized water. Colonies were counted manually and surviving fraction calculated as: colonies/(seeded colonies × plating efficiency). For cell viability assays, we used CellTiter-Glo Luminescent Cell Viability Assay (Promega) which measures total cellular ATP. 200 to 500 cells per well were seeded in white opaque 96-well plates. 24h later, cells were treated with the indicated concentrations of aph, checkpoint inhibitors, HU, PARP inhibitors for 6 days. Prior to measurement, 75μL of CellTiter-Glo reagent was added into each well and the contents were mixed for 30 min to induce cell lysis on an orbital shaker.

Luminescence was measured using the bioluminescence plate reader Envision 2102 (Perkin Elmer).

### Immunofluorescence staining

Anti-BrdU. To detect single stranded DNA under non-denaturing conditions, U2OS cells were seeded 24 h prior to treatment on 12 mm coverslips in 12-well plates. For parental ssDNA, BrdU (10 μM) was added for a total of 48h (24h before aph treatment and 24h with aph treatment). For nascent ssDNA, BrdU (20 μM) was added to aph-treated cells 10 min before and during the 4h treatment with checkpoint inhibitors. Cells were permeabilised at 4°C for 10 min in PBS + 0.5% Triton X-100 followed by fixation at room temperature (RT) for 10 min in 4% paraformaldehyde (PFA). Cells were fixed again at −20°C with 100% methanol for 10 min, dried for 1 min, and then washed with PBS. Cells were then incubated at RT with blocking solution -PBS-T and 2% BSA-for 1h. Coverslips were then incubated with primary antibodies: mouse anti-BrdU (BD Biosciences; clone B44, 1:200 at 37°C for 1h. Coverslips were washed for 5 min in PBS-T and incubated at RT for 1h with secondary antibodies (1:500; Alexa Fluor 488). Coverslips were then washed with PBS-T and stained with DAPI (1 μg/mL) and mounted on microscope slides with Fluoroshield (Sigma). **RPA and Rad51.** U2OS cells were seeded on 12 mm coverslips in 12-well plates 24 h prior to treatment. Cells were treated with the indicated drugs. Coverslips were then incubated in cytoskeleton buffer (CSK) 0.5% Triton X-100 for 5 min on ice and fixed with 4% PFA for 20 min at RT. Coverslips were then washed twice in PBS, permeabilized with 0.1% Triton X-100 in PBS for 15 min at 4°C and blocked in 5% BSA 0.1% Triton X-100 in PBS overnight. Primary incubation was performed overnight in blocking solution using the anti-RAD51 antibody (Abcam; ab133534; 1:100). Coverslips were then washed with PBS containing 0.25% Tween 20. Secondary incubation was performed in PBS using Alexa 488 (1:500) for 1h at RT. Coverslips were then washed with PBS-T and stained with DAPI (1 μg/mL) and mounted on microscope slides with Fluoroshield (Sigma).

### Neutral Comet Assay

Neutral comet assays were performed using a Comet Assay kit from Amsbio (4250-050-K) according to the manufacturer’s protocol.

### Cell fractionation

1×106 U2OS cells were seeded. 24h later cells were treated for 24h with aph alone or in combination with checkpoint inhibitors (AZD-7662, 100nM) before harvesting for the indicated times. Cells were scraped in 1ml ice-cold PBS and spun down for 4 min at 500 g and resuspended in 200ul CSK buffer (10mM PIPES pH7.0, 100mM NaCl, 300mM Sucrose, 3 mM MgCL2, 1mM EGTA, 0.5% Triton, 1mM DTT and 1x of protease inhibitor mix (Complete, EDTA-free, Roche). Samples were incubated on ice for 10 min. Cells were spun down for 3 min at 1000 rpm. 150ul of supernatant, representing the soluble fraction, was collected and 50ul of 4x Laemmli Buffer (Bio-Rad) + 10% 2-metcaptoethanol (Sigma-Aldrich) was added. Residual soluble fraction was removed, and the chromatin pellet was washed in 500ul of CSK. 200uL of 1x Laemmli Buffer + 2.5% 2-metcaptoethanol was added. Samples were sonicated with a probe at medium intensity (10A) for 12 s in a Branson 250 instrument, and then incubated at 95°C for 10 min. 20ul of each sample was loaded and subjected to SDS-PAGE and immunoblotting.

## Acknowledgments

We thank Jiri Lukas for the gift of the SuperRPA cell line. We are grateful to the high throughput screening, light microscopy and flow cytometry science technology platforms at the Francis Crick Institute. We thank Allison McClure for stimulating discussions.

This work has received funding from the European Union’s Horizon 2020 research and innovation programme under the Marie Skłodowska-Curie (895786 to BC), Wellcome Trust Senior Investigator Awards (106252/Z/14/Z and 219527/Z/19/Z to JFXD), and a European Research Council Advanced Grant (669424-CHROMOREP to JFXD). This work was supported by the Francis Crick Institute, which receives its core funding from Cancer Research UK (FC001066), the UK Medical Research Council (FC001066), and the Wellcome Trust (FC001066). For the purpose of Open Access, the author has applied a CC BY public copyright licence to any Author Accepted Manuscript version arising from this submission.

## Author contributions

Conceptualization: APB, BC, JFXD

Methodology: APB, BC, JFXD

Investigation: APB, BC, MY, JZ, RI, MH

Writing – original draft: APB, BC, JFXD

Writing – review & editing: APB, BC, JFXD

Supervision: JFXD

Project administration: APB, BC, JFXD

Funding acquisition: BC, JFXD

## Supplementary Figure legends

**Figure S1. Related to Figure 1 and Figure 2.**
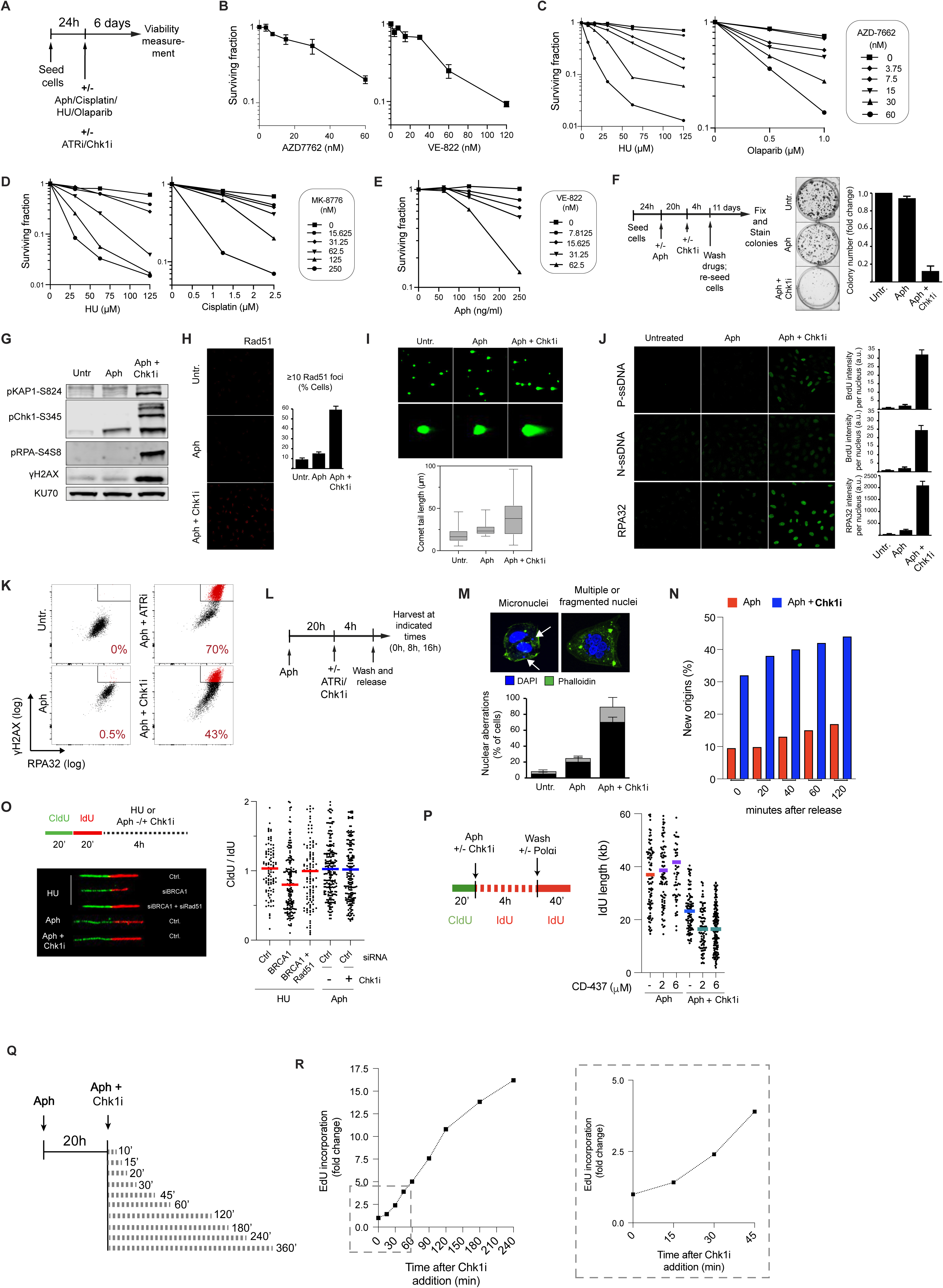
**(A)** Experimental setup for cell viability assays using CellTiter-Glo. **(B-E)** U2OS cells were treated as indicated in **Figure S1A** with checkpoint inhibitors (ATRi, VE-822; Chk1i, AZD7662 and MK-8776) alone or in combination with different genotoxic drugs at the indicated concentrations. Cell viability was normalised to cells untreated with genotoxic drugs for each checkpoint inhibitor condition. **(F)** Clonogenic survival of U2OS cells. Left panel, experimental set up. After drugs were washed off, cells were released in complete media and allowed to grow for 14 day. Colonies were then fixed and stained with crystal violet. Middle panel, representative images of colony staining. Right panel, quantification showing the fold change in colony number relative to untreated condition. Error bars represent s.e.m. **(G)** Western blot analysis of DNA damage markers. Following treatment, cells were immediately harvested. Ku70 protein levels were used as loading controls. **(H)** Immunofluorescence analysis of chromatin-bound Rad51. Upper panel, representative images. Bottom panel, graph showing percentage of cells with equal or more than 10 Rad51 foci. **(I)** Assessment of DSB formation by neutral comet assay. Upper panel, representative comets. Bottom panel, quantification of comet length. At least 50 comets were scored per sample. **(J)** Immunofluorescence analysis of parental and nascent single strand DNA (P-ssDNA and N-ssDNA) and chromatin-bound RPA32. Left, representative images. Right, quantifications. **(K)** Flow cytometry analysis of the levels of chromatin-bound RPA32 and γH2AX in the same population of cells. Cells were left untreated or treated with 1 μg/ml aphidicolin (aph) alone or in combination with checkpoint inhibitors (ATRi, VE-822; Chk1i, AZD7662). Representative images comparing γH2AX intensity (log scale) and chromatin-bound RPA32 intensity (log scale) in cells. Gated cells depicted in red represent cells with simultaneously high levels of γH2AX and RPA. Bottom right of each panel, quantification of the percentage of red coloured cells. **(L)** Experimental setup used for Figures 1B, **1C** and **S1M**. Cells were seeded and after 24h left untreated or treated with 1 μg/ml aphidicolin (aph) alone or in combination with 50nM Chk1i (AZD7662). Following treatment, cells were immediately harvested, or drugs were washed, and cells were released in complete media which depending on the experiment contained no drug, 100 nM taxol or 4.5 μg/ml cytochalasin B. **(M)** Genomic instability analysis. U2OS cells were treated like Figure 1L but analysed forty hours after release in complete media with 4.5 μg/ml cytochalasin B. Cells then were fixed, stained with DAPI and phalloidin and analyzed for nuclear aberrations. Upper panel, representative images. Bottom panel, graph showing percentage of cells showing nuclear aberrations. **(N)** Percentage of new origins fired in samples shown in Figures 1D-**1F** was determined by counting the number of IdU/red-only fibers over the total number of different replication structures. At least 600 of total structures were scored per sample. **(O)** Experimental setup to assess nascent DNA resection: cells were pulsed with CldU for 20 min, washed, then incubated with IdU for 20 min, washed and treated with HU (4 mM) or 1 μg/ml aph alone or in combination with 50 nM Chk1i for 4h. Left, representative micrographs of CldU and IdU replication tracks. Right, scatter plot and mean IdU track length in bicolour fibers. At least 150 forks were scored per sample. **(P)** DNA fiber analysis of replication fork progression. Left, Experimental setup. Cells were pulsed with CldU for 20 min, washed and then incubated with IdU plus 1 μg/ml aph alone or in combination with 50 nM Chk1i for 4h. Cells were then washed and released in complete media containing IdU and the Polα inhibitor CD-437 for 40 min. Right, scatter plot and mean IdU track length in bicolour fibers is shown. At least 150 forks were scored per sample. **(Q)** Timing of events following checkpoint inhibition. Schematic of the experimental approach used in Figures 2A, **2B** and **S2B**. U2OS cells were treated with 1 μg/ml aph for 20h and then Chk1i (AZD7662, 50 nM) was added for the indicated times. **(R)** EdU incorporation of cells treated with Chk1i at different times. Graphs show fold change of mean fluorescence intensities (MFI) of EdU of each time point in comparison with aph treatment alone. Right, graph inset for 0-45min of Chk1i treatment.

**Figure S2. Related to Figure 3.**
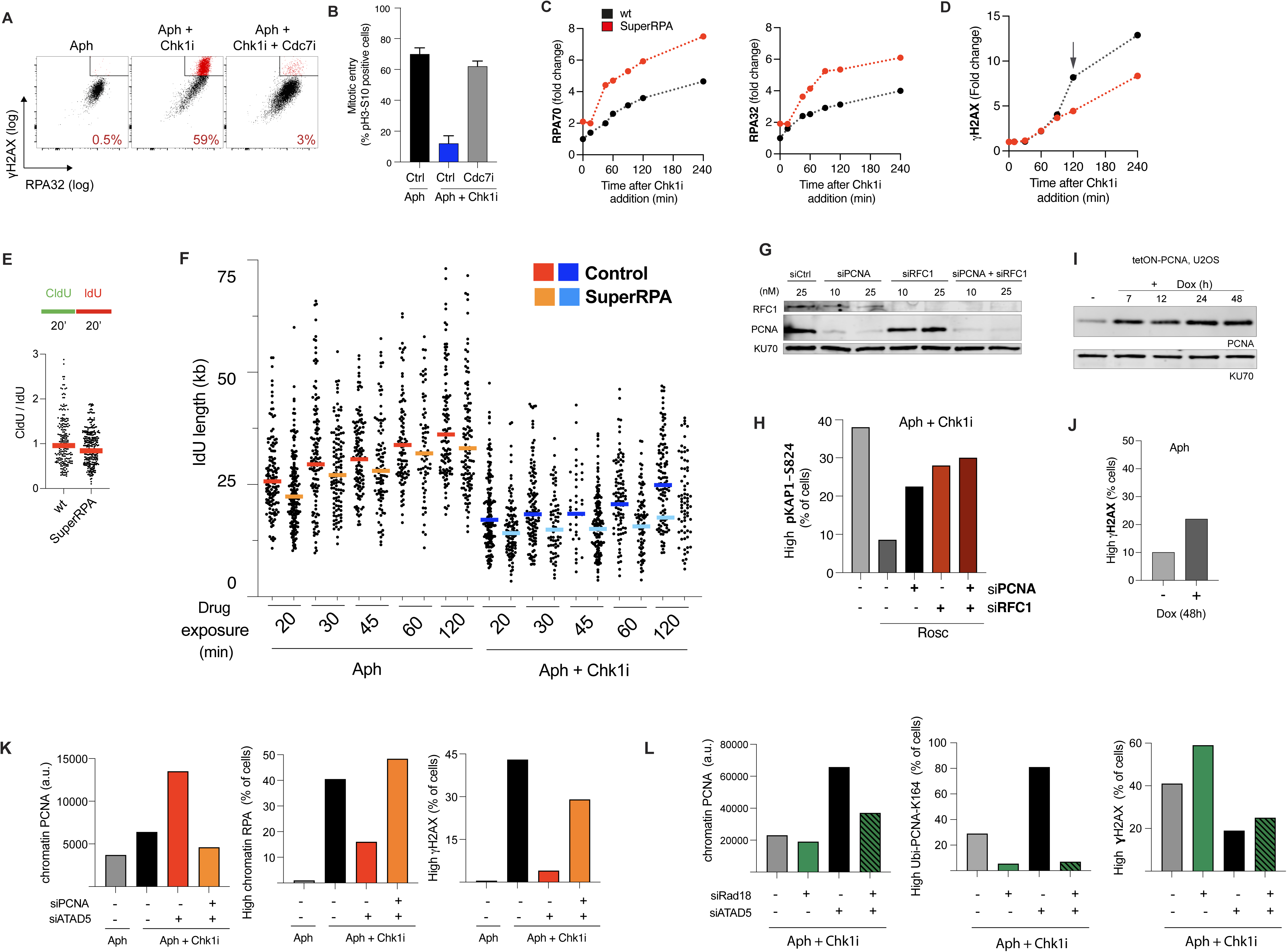
**(A, B)** Inhibition of new origin firing rescues checkpoint inhibition phenotypes. In experiments shown in **Figures S3A**, **S3B** and **3A**-**3C**, cells were pre-treated with Cdc7i (XL143, 10 μM) or CDK2i (roscovitine, 10 μM) for 60 min, and then incubated in combination with the checkpoint inhibitor (AZD7662). **(A)** Flow cytometry analysis of chromatin-bound RPA32 and γH2AX levels in the same population of cells. Cells were treated with aph alone and aph/Chk1i alone or in combination with CDK2i (10 μM). Representative images comparing γH2AX intensity (log scale) and chromatin-bound RPA32 intensity (log scale) in treated cells. Gated cells depicted in red represent cells with simultaneously high levels of γH2AX and RPA. Right bottom corner, quantification of the percentage of red coloured cells in each sample. **(B)** Mitotic entry analysis. Cells were treated like **Figure S1L** and 16 hours after release in complete media with 100 nM taxol, cells were fixed and stained for phosphoH3-S10 and DAPI followed by flow cytometric analysis. The proportion of pH3-S10 positive cells in each sample is shown. **(C-F)** Overexpression of RPA complex using the SuperRPA cell line does not rescue early phenotypes of checkpoint inhibition. Flow cytometry analysis of chromatin-bound RPA32, RPA70 **(C)** and γH2AX **(D)** in wild-type and SuperRPA cells treated with aph and Chk1i for different times. Graphs show fold change of mean fluorescence intensities (MFI) of each protein for each time point in comparison with aph treatment alone.**(E)** DNA fiber analysis of replication fork progression in unperturbed conditions in wild-type and SuperRPA cells. Cells were pulsed with CldU for 20 min, washed, then incubated with IdU for 20 min. Scatter plot and mean IdU track length in bicolour fibers is shown. At least 200 forks were scored per sample. **(F)** DNA fiber analysis of replication fork progression in wild-type and SuperRPA cells. The experimental set up used is depicted in Figure 2C. Cells were pulsed with CldU for 20 min, washed and then incubated with IdU plus 1 μg/ml aph alone or in combination with 50 nM Chk1i for the indicated amount of time. Cells were then washed and released in complete media containing IdU for 40 min. Scatter plot and mean IdU track length in bicolour fibers is shown. At least 200 forks were scored per sample. **(G)** Immunoblots of U2OS cells treated with PCNA siRNA, RFC1 siRNA or a combination of both in unperturbed conditions. Ku70 protein levels were used as loading controls. **(H)** Flow cytometry analysis of cells presenting high levels of pKAP1-S824 staining. Cells were treated with aph in combination with Chk1i and roscovitine depleted for PCNA and RFC1 alone or in combination. **(I)** Immunoblots of whole-cell extracts of tetON-PCNA U2OS cells showing PCNA expression after 7, 24, 48h of doxycycline (Dox) treatment. Ku70 protein levels were used as loading controls. **(J)** Flow cytometry analysis of the percentage of cells presenting high levels of γH2AX staining. Control cells and tetON-PCNA cells were left untreated or treated with 1 μg/ml aph for 48h. **(K)** Flow cytometry analysis of the chromatin-bound PCNA, chromatin-bound RPA and γH2AX levels. Control cells were treated with aph alone and control cells and cells depleted for ATAD5 alone or in combination with PCNA were treated with 1 μg/ml aph for 24h, with the addition of 50 nM Chk1 inhibitor during the final 4 hours. **(L)** Flow cytometry analysis of the chromatin-bound PCNA, ubiPCNA-K164 and γH2AX levels. Control cells and cells depleted for ATAD5 alone or in combination with Rad18 were treated with 1 μg/ml aph for 24h, with the addition of 50 nM Chk1 inhibitor during the final 4 hours.

**Figure S3. Related to Figure 4.**
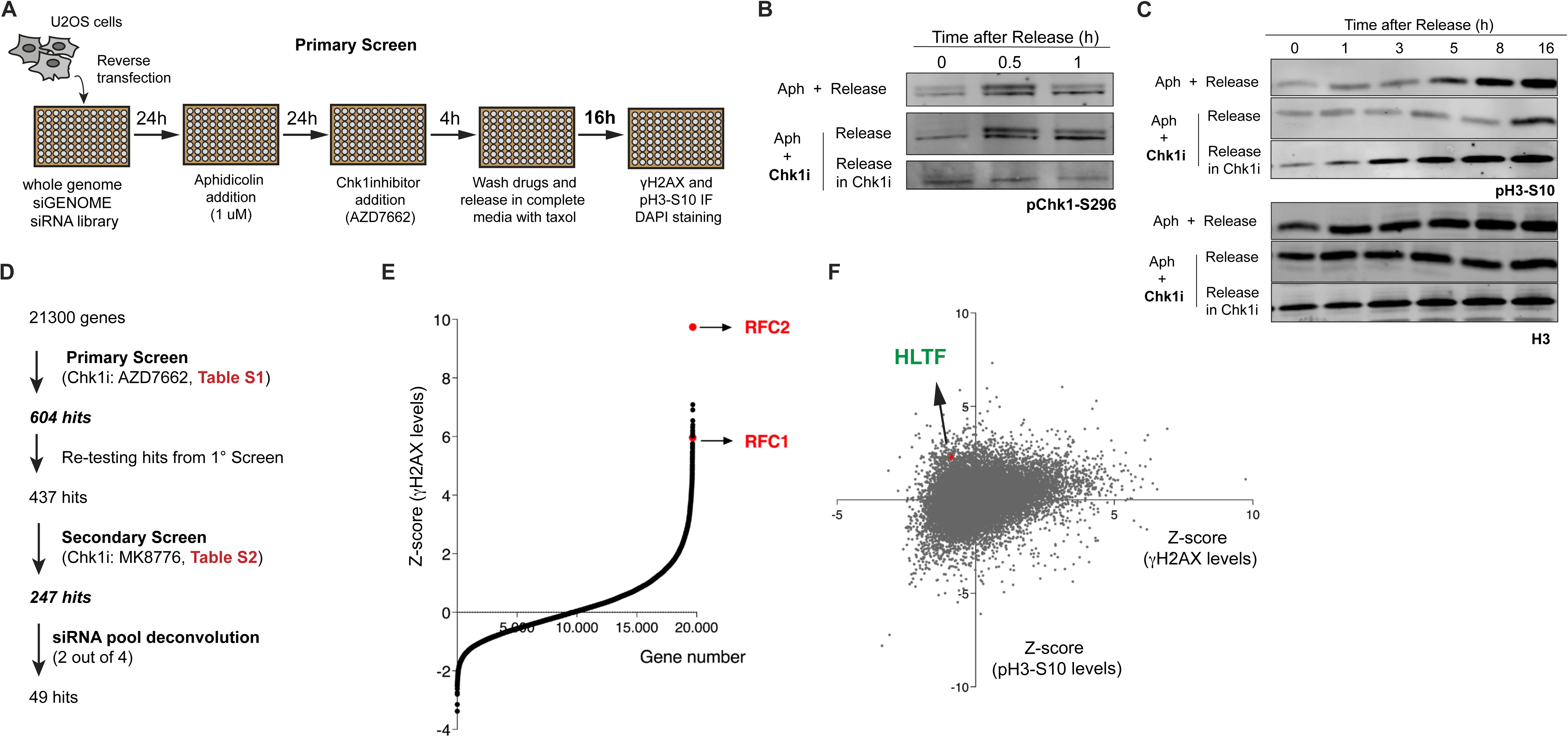
**(A)** Overview of the RNA interference-based genome-wide screen methodology. U2OS cells were plated in 96 well plates and reversed transfected with siRNAs. Twenty-four hours later, cells were treated with aph for 24h and then a reversible Chk1 inhibitor (50nM, AZD7762) was added for 4h. Aphidicolin and Chk1 inhibitor were then removed, and cells were released in medium containing taxol for 16h. Cells were harvested, stained for γH2AX and pH3-S10 and fluorescence intensity values of both markers were determined by fluorescence microscopy on an automated platform. **(B)** Chk1 reactivation after inhibition with Chk1 inhibitor AZD7662 occurs at around 30 min after inhibitor wash out. Immunoblot of U2OS cells showing phosphorylation of Chk1-S296 (Chk1 auto-phosphorylation). Cells were treated with 1 μg/ml aph for 24h, with the addition of 50 nM Chk1 inhibitor during the final 4 hours. Drugs were then washed out and cells were released in complete media. Samples were analysed before and after 30 and 60 min of release. **(C)** Chk1 reactivation after inhibition with Chk1 inhibitor AZD7662 triggers G2/M arrest. Immunoblot of U2OS cells showing phosphorylation of H3-S10 (mitotic marker). Cells were treated with 1 μg/ml aph for 24h, with the addition of 50 nM Chk1 inhibitor during the final 4 hours. Drugs were then washed out and cells were released in complete media with 100 nM of taxol for 1, 3, 5, 8 and 16 hours. **(D)** Graphical summary of the study workflow from the successive siRNAs screens. The primary screen was performed independently two times and identified 437 overlapping siRNAs that reduced γH2AX and increased pH3-S10 levels by at least 1.5 standard deviation from the population mean (**Table S1**). Those 437 siRNAs were re-tested in a secondary screen, with the same conditions as the primary screen but with a different Chk1 inhibitor (MK-8776), yielding a total of 247 siRNAs (**Table S2**). siRNAs were considered for further studies when simultaneously showed a decrease in γH2AX levels indicating reduced DNA damage and an increase pH3-S10 levels indicating higher levels of mitotic entry. *This criterion was chosen to detect siRNAs that allow a ‘functional’ rescue of the irreversibility of the replication arrest as not only DNA damage was monitored but also, and most importantly, the ability of these forks to resume replication, complete S phase and enter into mitosis.* Next, the 121 top scoring siRNA pools were re-screened by siRNA deconvolution. For each siRNA pool chosen, the four siRNAs comprising the original pool were individually tested for increased pH3-S10 levels in the same conditions. Forty-nine genes gave rise to a clear increased mitotic index (≥ 30% compared to control) with at least 2 out of 4 siRNA oligos. **(E)** Primary screen z-score distribution of all siRNAs analysed for DNA damage assessed by γH2AX levels. Each dot represents the average Z score of an individual siRNA tested in triplicate. Out of the 21,000 genes tested, RFC1 depletion ranked among the top 20, while RFC2 depletion emerged as the highest-scoring gene. **(F)** Primary screen z-scores distribution of all siRNAs analysed for DNA damage assessed by γH2AX levels and for mitotic entry assessed by pH3-S10. One of the strongest hits we recovered for both parameters was helicase-like transcription factor (HLTF), a RING domain-containing ubiquitin ligase and SNF2-family DNA translocase previously implicated in several DNA repair/tolerance pathways.

**Figure S4. Related to Figure 4.**
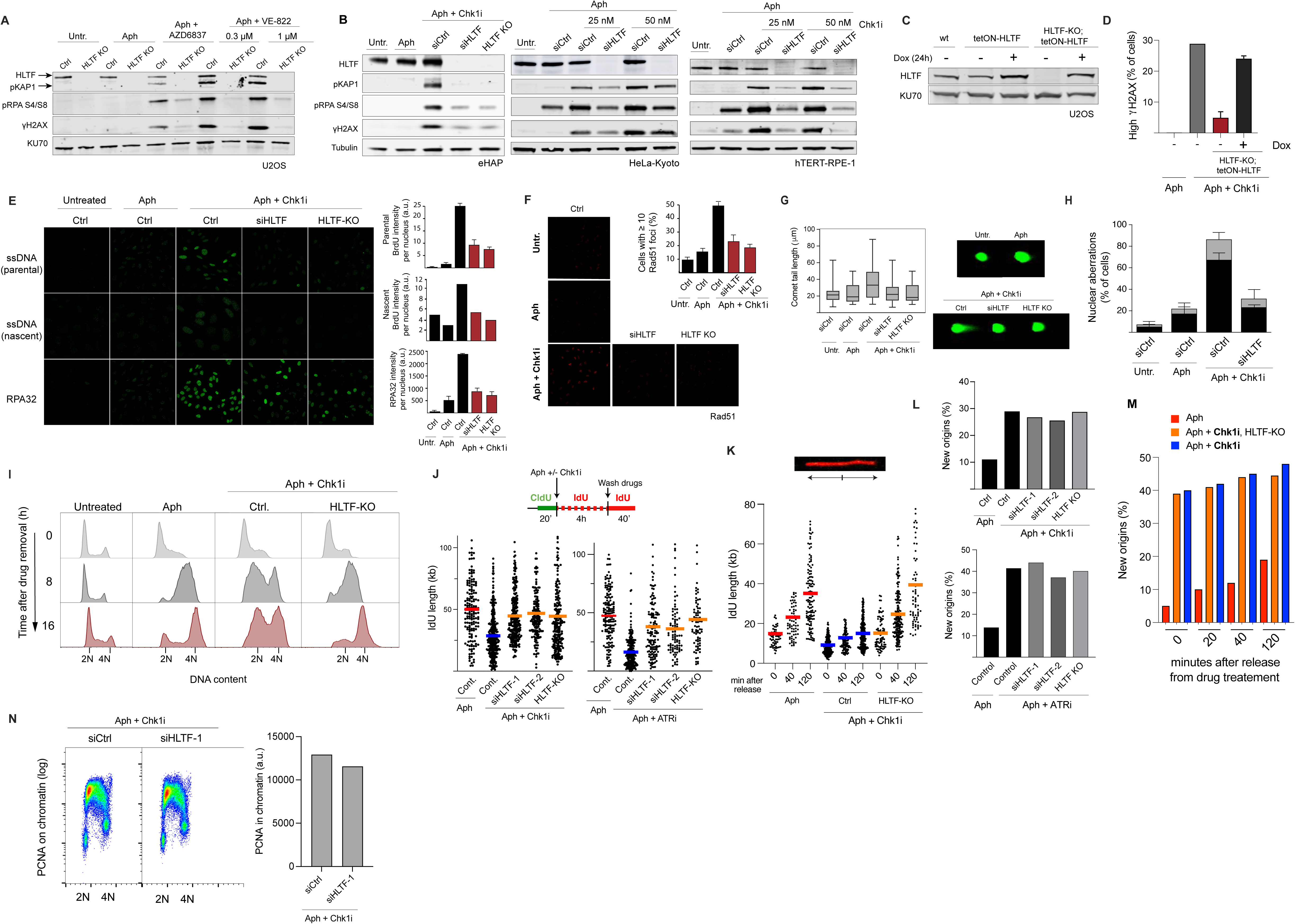
**(A)** Immunoblots showing of DNA damage markers (pKAP1, pRPA-S4S8 and γH2AX) in control and HLTF CRISPR knock out (KO) U2OS cells after Chk1 and ATR inhibition. Control and HLTF KO cells were left untreated, treated with 1 μg/ml aph for 24h alone or during the final 4 hours in combination with 50 nM Chk1 inhibitor. After treatment, cells were immediately harvested. Ku70 protein levels were used as loading controls. **(B)** Immunoblots showing of DNA damage markers (pKAP1, pRPA-S4S8 and γH2AX) after Chk1 inhibition in control HLTF KD in eHAP, HeLa and RPE1 cells and CRISPR KO eHAP cells. Control cells were left untreated or treated with 1 μg/ml aph for 24h alone. Control and HLTF KO/KD cells were treated with aph for 24h and Chk1i was added during the final 4 hrs. After treatment, cells were immediately harvested. Tubulin levels were used as loading controls. **(C)** Immunoblots showing HLTF expression by doxycycline (Dox) treatment in tetON-HLTF generated in a wild type U2OS background (left) or HLTF-KO background (right). **(D)** Flow cytometry analysis of the percentage of cells presenting high levels of γH2AX staining. Control cells were treated with 1 μg/ml aph for 24h alone and control, HLTF KO cells and HLTF-KO/tetON-HLTF were treated with aph for 24h and a checkpoint inhibitor was added during the final 4 hrs. After treatment, cells were immediately harvested. **(E)** Immunofluorescence analysis of parental and nascent single strand DNA, chromatin-bound RPA32 in control, HLTF knock down and HLTF KO cells treated like **Figure S5A**. Left, representative images. Right quantification of BrdU or RPA32 intensity. **(F)** Immunofluorescence analysis of chromatin-bound Rad51 in HLTF-KD and HLTF-KO cells after checkpoint inhibitor treatment. Right, representative images. Left, graph showing percentage of cells with equal or more than 10 Rad51 foci. **(G)** Assessment of DSB formation by neutral comet assay. Left, quantification of comet length. Right, representative comets. At least 40 comets were scored per sample. **(H)** Genomic instability in control and HLTF deficient cells. Control cells were left untreated or treated with 1 μg/ml aph for 24h alone. Control and HLTF KD cells were treated with aph for 24h and a checkpoint inhibitor was added during the final 4 hrs. Forty hours after release in complete media containing 4.5 μg/ml cytochalasin B, cells were fixed, stained with DAPI and phalloidin and analyzed for nuclear aberrations. Graph shows percentage of cells showing nuclear aberrations. **(I)** Flow cytometry analysis of S phase progression. Control cells were left untreated or treated with 1 μg/ml aph for 24h alone. Control and HLTF KO cells were treated with aph for 24h and a checkpoint inhibitor was added during the final 4 hrs. 0h, 8h and 16h after release in complete media with 100 nM taxol, cells were fixed and stained with DAPI followed by flow cytometric analysis. Cell cycle histograms are shown. **(J-L)** DNA fiber analysis of replication fork progression and origin firing in HTLF-KO and HLTF-KD U2OS cells. (**J**) Upper panel, experimental set up: cells were pulsed with CldU for 20 min, washed and then incubated with IdU plus 1 μg/ml aph alone or in combination with Chk1i (AZD7662, 50 nM) or ATRi (VE-822, 500 nM) for 4h. Cells were then washed and incubated again only with IdU for 40 min. Bottom panel, scatter plot and mean IdU track length in bicolour fibers. At least 150 forks were scored per sample. **(K)** Scatter plot and mean IdU track length in red-only fibers from samples shown in Figures 4D and **4E**. **(L)** Percentage of new origins fired by DNA fiber analysis in samples shown in **Figure S5J**. **(M)** Percentage of new origins fired by DNA fiber analysis in samples shown in Figures 4D and **4E**. The percentage of new origins fired during checkpoint inhibition was determined by counting the number of IdU/red-only fibers over the total number of different replication structures. At least 600 of total structures were scored per sample. **(N)** Flow cytometry analysis of chromatin-bound PCNA levels. Control cells or HLTF siRNA cells were treated with aph and Chk1i as described in **Figure S5A**.

**Figure S5. Related to Figure 4.**
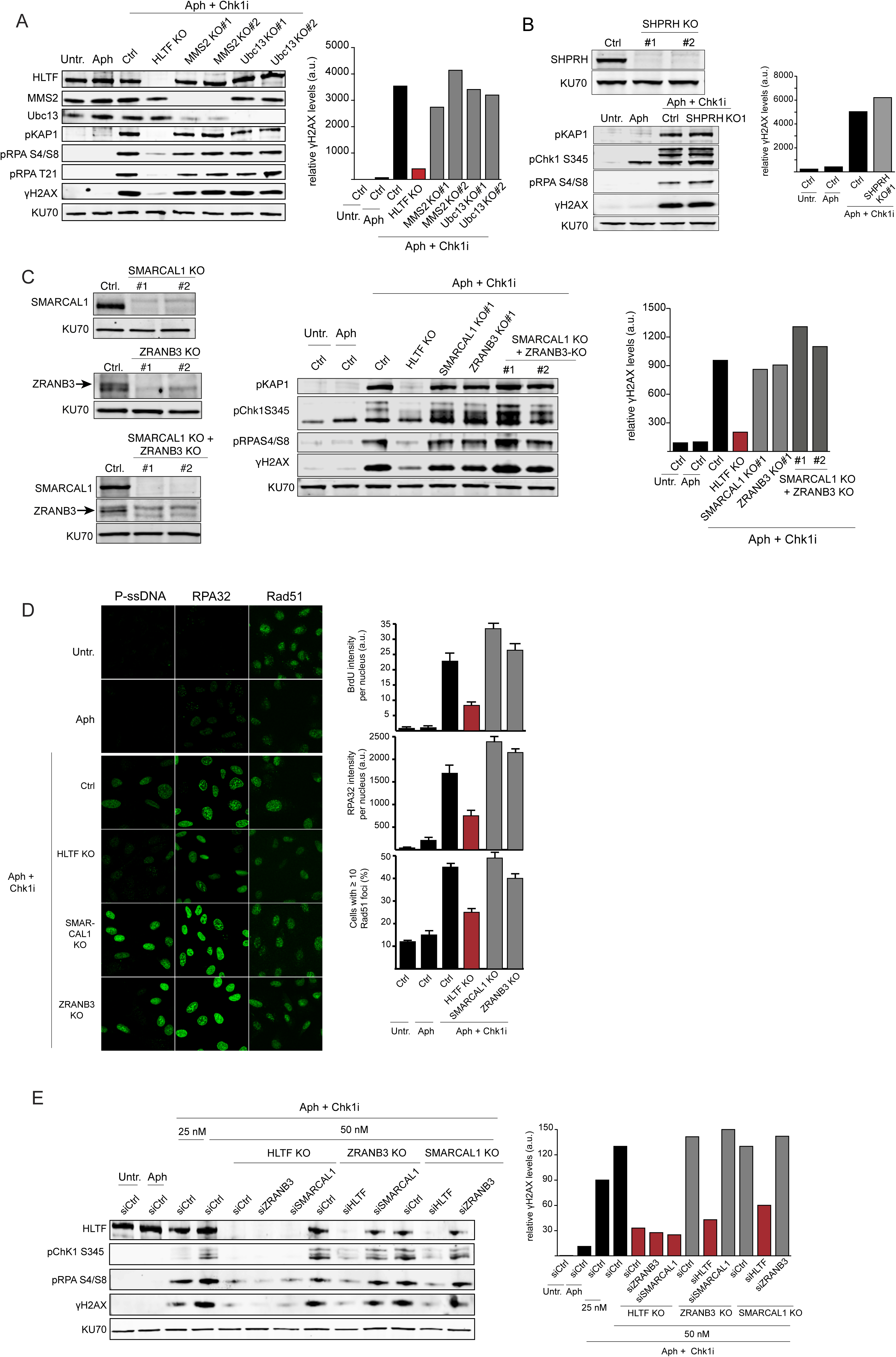
**(A)** MM2/Ubc13 PCNA, heterodimeric E2 ubiquitin conjugatin enzymes involved in the PCNA polyubiquitylation pathway, were individually knocked out in U2OS cells. DNA damage was monitored by immunoblotting of pKAP1, pRPA-S4/S8, pRPA-T21 and γH2AX, in cells treated with aphidicolin +/− Chk1 inhibitor. Right panel, quantification of γH2AX intensity in the blot (as images were acquired with a LICOR instrument in the linear range for quantification purposes). **(B)** SHPRH, a human orthologe of yeast Rad5 like HLTF, was individually knocked out in U2OS cells. DNA damage was monitored by immunoblotting of pKAP1, pCHK1-S345, pRPA-S4/S8, and γH2AX in cells treated with aphidicolin +/− Chk1 inhibitor. Right panel, quantification of γH2AX intensity in the blot. **(C)** ZRANB3 and SMARCAL1 were knocked out in U2OS cells, individually or in combination. DNA damage was monitored by immunoblotting of pKAP1, pCHK1-S345, pRPA-S4/S8, and γH2AX in cells treated with aphidicolin +/− Chk1 inhibitor. Right panel, quantification of γH2AX intensity in the blot. **(D)** In ZRANB3 or SMARCAL1 individual knockout cells treated with aphidicolin, with or without Chk1 inhibitor, DNA damage was assesed by immunofluorescence analysis of parental single strand DNA (P-ssDNA), chromatin-bound RPA32 and chromatin-bound Rad51. Left panel, representative images. Right panel, quantifications. **(E)** HLTF, ZRANB3, and SMARCAL1 knockout cells were treated either alone or in combination with siRNAs targeting HTLF, ZRANB3, or SMARCAL1, as indicated. DNA damage was monitored by immunoblotting of pCHK1-S345, pRPA-S4/S8, and γH2AX in cells treated with aphidicolin +/− Chk1 inhibitor. Right panel, quantification of γH2AX intensity in the blot.

**Figure S6. Related to Figure 4.**
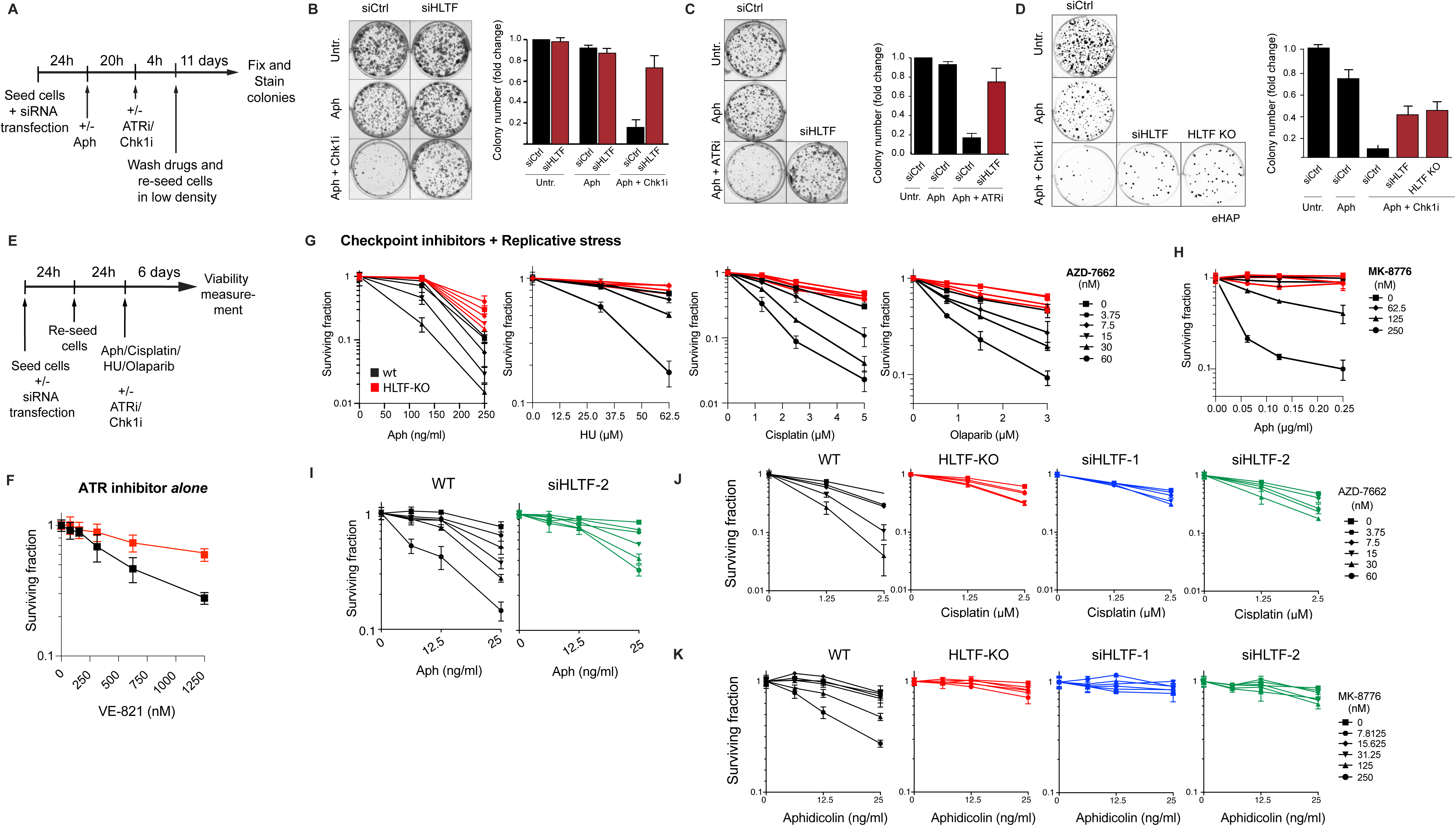
**(A-D)** Clonogenic survival of control and HLTF siRNA treated cells. **(A)** Experimental setup. **(B-C)** U2OS cells were treated with Chk1i (AZD7662, 50 nM) (**B**) or ATRi (VE-822, 500 nM) (**C**). **(D)** HLTF-KD and HLTF-KO eHAP cells were treated with Chk1i (AZD7662, 50 nM). Left, representative images of colony staining. Right, graph shows the fold change in colony number relative to the untreated condition. Error bars represent s.e.m. **(E-K)** Cell viability assays of HLTF-KD and HLTF-KO U2OS cells using CellTiter-Glo.**(E)** Experimental setup. **(F)** Survival assay of HLTF-KO cells treated with indicated concentrations of ATRi alone (VE-821). **(G, H)** HLTF-KO U2OS cells were treated as indicated in **Figure S6E** with the specified concentrations of Chk1i (AZD7662, **G** or MK-8776, **H**) alone or in combination with different genotoxic drugs at the indicated concentrations. Cell viability was normalised to cells untreated with genotoxic drugs for each checkpoint inhibitor condition. **(I-K)** HLTF-KO U2OS cells and cells treated with two different HLTF siRNAs were subjected to the treatment shown in **Figure S6E** with the indicated concentrations of Chk1i (AZD7662, **I** and **J** or MK-8776, K) alone or in combination with genotoxic drugs at the indicated concentrations (aph, **I** and **K** or cisplatin, **J**). Cell viability was normalised to cells untreated with genotoxic drugs for each checkpoint inhibitor condition.

